# Pooled effector library screening in protoplasts rapidly identifies novel *Avr* genes

**DOI:** 10.1101/2023.04.28.538616

**Authors:** Taj Arndell, Jian Chen, Jana Sperschneider, Narayana M Upadhyaya, Cheryl Blundell, Nathalie Niesner, Aihua Wang, Steve Swain, Ming Luo, Michael A Ayliffe, Melania Figueroa, Thomas Vanhercke, Peter N Dodds

## Abstract

Crop breeding for durable disease resistance is challenging due to the rapid evolution of pathogen virulence. While progress in resistance (*R*) gene cloning and stacking has accelerated in recent years ^1-3^, the identification of corresponding avirulence (*Avr*) genes in many pathogens is hampered by the lack of high-throughput screening options. To address this technology gap, we developed a platform for pooled library screening in plant protoplasts for rapid identification of interacting *R*/*Avr* pairs. We validated this platform by isolating known and novel *Avr* genes from wheat stem rust (*Puccinia graminis* f. sp. *tritici*) by screening a designed library of putative effectors against individual *R* genes. Rapid *Avr* gene identification provides molecular tools to understand and track pathogen virulence evolution by genotype surveillance and optimise *R* gene deployment and stacking strategies. This screening platform is broadly applicable to many crop pathogens, whilst also adaptable for screening genes involved in other protoplast-selectable traits.

---

Crop pathogens greatly reduce agricultural productivity and are a persistent threat to global food security ^4, 5^. The most effective and sustainable approach to mitigate crop disease is through breeding of resistance (*R*) genes into crop varieties. Most *R* genes encode immune receptors, such as nucleotide-binding leucine-rich repeat (NLR) proteins, that directly or indirectly recognise pathogen effectors, known as avirulence (Avr) proteins ^6, 7^. These immune recognition events induce plant defense responses, often including localised cell death, which limit pathogen spread. However, pathogens continuously evolve to escape recognition via mutation of *Avr* genes. Thus, understanding *Avr* gene diversity in pathogen populations is critical for the effective deployment of *R* genes in breeding or gene stacking. However, the complex genomes of filamentous pathogens, often encoding hundreds to thousands of effector candidates ^8^, pose challenges for *Avr* gene identification.

Rust fungi (order Pucciniales) cause serious diseases of many crop plants, especially among staple cereal crops including wheat, barley, oat and corn ^9^. For example, wheat stem rust disease is caused by the fungus *Puccinia graminis* f. sp. *tritici* (*Pgt*) and newly arisen virulent strains, such as Ug99, have caused devastating losses and threaten global wheat production. ^10, 11^. Rust fungi have large genomes with two separate haploid nuclei and encode thousands of potential effector genes that typically share minimal or no sequence homology to other proteins with known functions ^9^. Although hundreds of rust resistance loci are described in cereals (many no longer effective), only three corresponding *Avr* genes (*AvrSr27, AvrSr35, AvrSr50*) ^12-14^ have been identified in *Pgt* and only two from other cereal rusts; *AvrRppC* and *AvrRppK* from common corn rust *P. sorghi* ^15, 16^.

Approaches to functionally test *Avr* gene candidates usually involve pairwise transient co-expression of individual *R* and *Avr* gene combinations by PEG-mediated protoplast transformation or leaf agroinfiltration to detect immunity-induced cell death ^17, 18^. However, these methods assay candidate effectors one-by-one, so screening many effectors requires highly labour-intensive sequential assays ^14, 15^. The increasing availability of high-quality pathogen genome sequences combined with recent advances in fungal and oomycete effector prediction^8^ presents an opportunity to design and synthesise effector libraries to systematically screen genome-wide pathogen effector complements for immuno-recognition. We therefore set out to develop a platform for pooled effector library screening in plant protoplasts to enable rapid identification of interacting *R*-*Avr* combinations. Fig.1 outlines the selection scheme in which an *R* gene of interest and a pooled effector gene library are co-delivered to protoplasts such that a sub-population of cells expressing the *Avr* gene would undergo cell death, allowing identification based on the depletion of *Avr* gene transcripts in the living cell population.

**Fig. 1.**
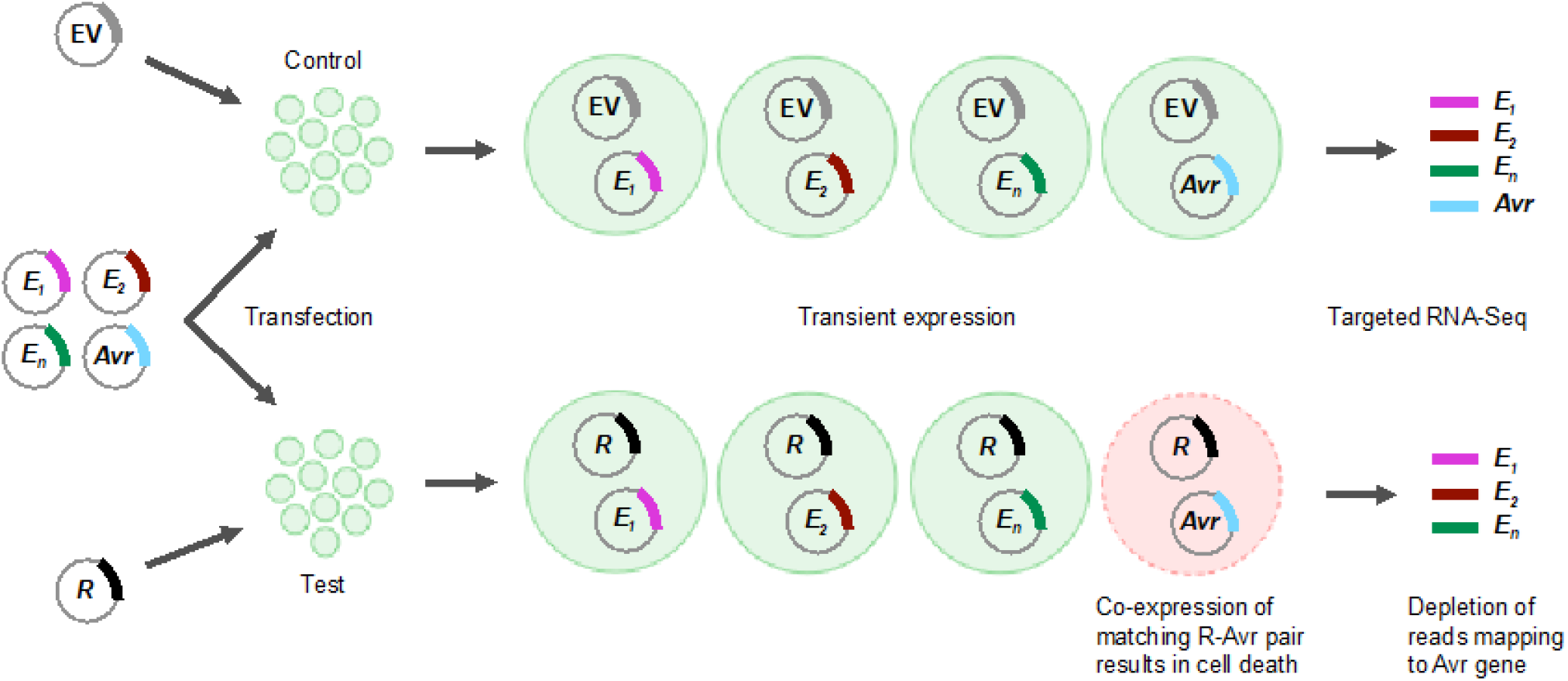
Schematic of effector library screening process to detect *Avr* genes. Protoplast populations are co-transformed with a pooled library of effector genes (E_1_, E_2_, etc.) from a pathogen along with either an empty vector (EV) for the negative control or a known *R* gene from a host plant. The library multiplicity of transfection (MOT), defined as the number of library plasmid molecules per cell, is chosen such that each protoplast individually receives a random but limited number of different effector candidates from the library together with either the empty vector or *R* gene. In the presence of the host *R* gene, protoplasts that express a matching *Avr* effector gene undergo programmed cell death, while cells expressing the same *Avr* effector gene in the negative control remain alive. Living protoplasts are subsequently collected from both transformed populations and subjected to targeted (library-specific) RNA-Seq. The expression of each effector in the library is then compared between the two samples. *Avr* gene candidates are identified by a depletion in derived transcripts in the sample expressing the corresponding *R* gene.

A key requirement for library screening is that individual cells independently express different library constructs to allow for differential screening. However, it has been suggested that this may not be possible in protoplasts due to the large amounts of plasmid typically used in protoplast transfections ^19^. We first addressed this limitation by using flow cytometry to test for independent transformation of protoplasts with two reporter genes. Constructs encoding yellow or red fluorescent protein (YFP, RFP) were co-delivered to protoplasts at various concentrations, defined as the multiplicity of transfection (MOT) in million molecules per cell (Fig. 2a, Supplementary Fig. 1). At high MOTs (36, 18, 3.6 M molecules/cell) a high proportion of protoplast cells expressed both fluorescent markers (24-40%) with only a small fraction expressing a single marker. However, the proportion of doubly transformed cells dropped at lower MOTs, with only ∼1% of cells expressing both markers when delivered at 0.07 M mol/cell, while a larger proportion expressed a single marker (2-3% for each). This indicated that differential library screening may be feasible when individual constructs are delivered at these levels.

**Fig. 2.**
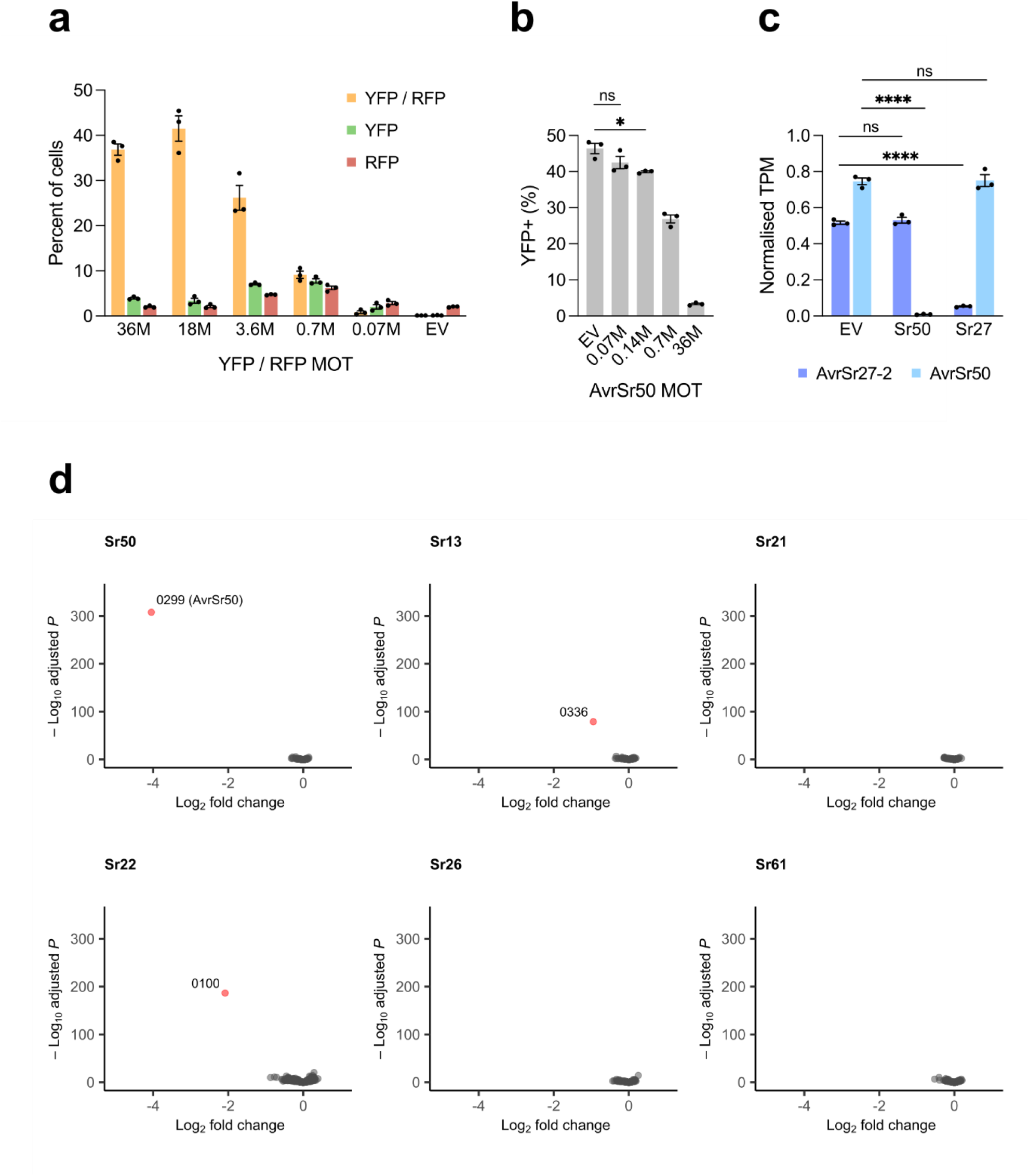
Protoplast library screen for R-Avr interactions. **a**, Protoplasts were co-transformed with YFP and RFP at different MOTs (million plasmid molecules per cell) and the percentage of living cells showing fluorescence in the YFP, RFP or both wavelengths was determined. Mean of three replicates with standard error shown. **b**, Protoplasts were co-transformed with YFP and *Sr50* plus a mock library consisting of *AvrSr50* at various MOTs within a background of *AvrSr35* (combined MOT 36). Plot shows the percentage of YFP-positive living cells (mean of three replicates [grey dots] with significant differences relative to the empty vector (EV) indicated (*, *p* < 0.05; two-sample *t*-test equal variances). **c**, Expression levels of AvrSr27-2 and AvrSr50 in protoplasts co-transformed with a mock library consisting of *AvrSr50* (MOT 0.14) and *AvrSr27-2* (MOT 0.14) within a background of *AvrSr35* (combined MOT 100), and either *Sr50, Sr27*, or an empty vector (EV). Data is the mean of three replicates and error bars represent the standard deviation. Significant differences are indicated (****, *p* < 0.0001). **d**, Expression analysis of a pooled stem rust effector library (dots) co-transformed into wheat protoplasts with wheat resistance genes *Sr50, Sr13c, Sr21, Sr22, Sr26* and *Sr61* compared to the empty vector. Graphs show volcano plots of differential expression (X-axis) versus adjusted P value (Y-axis). Effectors showing significantly reduced expression in each treatment are labelled.

Another key parameter for the screening approach is the induction of cell death in a sub-population of cells after library transformation. Previous assays for immunity-induced cell death in protoplasts have relied on co-expression of a luciferase reporter gene with corresponding *R* and *Avr* genes, with recognition resulting in reduced luciferase activity in the cell suspension as a whole relative to a control ^3, 18, 20^. We modified this approach to develop an individual cell scoring assay in which a fluorescent protein reporter (YFP) is used instead of luciferase, and transformed protoplasts are analysed individually for fluorescence by flow cytometry. Co-expression of three known wheat stem rust *R*-*Avr* pairs, *Sr50*-*AvrSr50, Sr27*-*AvrSr27-2*, and *Sr35*-*AvrSr35*, resulted in a significant decrease in the proportion of YFP-positive protoplasts in the living (propidium iodide-negative) population of cells, compared to the controls where single *R* or *Avr* genes or non-matching *R-Avr* pairs were expressed (Supplementary Fig. 2). This assay allows for quantification of the proportion of cells showing immune-induced cell death in a protoplast suspension. To simulate library screening conditions and determine whether cell death responses could still be recorded at low MOTs, we delivered *AvrSr50* at various MOTs within a background of *AvrSr35* (combined MOT 36M) along with *Sr50*. We observed a small but significant reduction in the proportion of YFP-positive cells with *AvrSr50* at MOT 0.14M (Fig. 2b), indicating that a subpopulation of cells expresses *AvrSr50* and undergoes cell death.

As a final validation step for the library screening platform, we tested whether differential effector gene expression could be used for identifying *Avr* gene candidates. For this, we transformed protoplasts with a mock library consisting of three constructs encoding *AvrSr50, AvrSr27-2* (each delivered at MOT 0.14 M) and *AvrSr35* (at MOT 100 M) along with either *Sr50, Sr27*, or an empty vector (at MOT 36 M). RNA-Seq analysis of *Avr* gene expression in protoplasts 24 hours post-transfection (Figure 2c) showed that both *AvrSr27-2* and *AvrSr50* were both expressed when the mock library was co-transformed with an empty vector. However, expression of *AvrSr50* was substantially reduced when co-expressed with *Sr50* but not with *Sr27*, and *AvrSr27-2* was reduced when co-expressed with *Sr27* but not with *Sr50*. This clearly indicated that the effects of each *Avr* gene could be independently assessed by their relative expression when co-delivered as part of a mock library.

Having established and optimised the experimental conditions for library screening, we synthesised a library consisting of 696 predicted *Pgt* effectors selected from the Pgt21-0 reference genome annotation ^11^ as genes encoding secreted proteins with expression patterns similar to known *Avr* genes ^12^. This library was pooled and co-transformed (MOT 0.14M per construct) into protoplasts with either an empty vector or one of seven separate *R* gene constructs encoding *Sr50, Sr27, Sr13c, Sr21, Sr22, Sr26*, or *Sr61*. RNA-Seq analysis was used to identify effectors showing reduced expression when co-expressed with specific *R* genes relative to the empty vector. Two independent screens correctly identified *AvrSr50* as a single gene showing significantly reduced expression in the presence of *Sr50* only (Fig. 2d, Supplementary Fig 3). Likewise, the five known variants of *AvrSr27* ^12, 21^ all showed reduced expression only when co-expressed with *Sr27* (Supplementary Fig. 3). Thus, this platform can specifically identify single and multiple *Avr* genes from a complex library of effector candidates screened against different *R* genes in parallel. Two effectors (library ID’s 0336 and 0100) showed significantly reduced expression only in the presence of *Sr13c* or *Sr22*, respectively, and represent candidates for *AvrSr13* and *AvrSr22*. Both candidates were identified in the two independent screens, highlighting the robustness of the platform. No effectors showed reduced expression in the presence of *Sr21, Sr26* or *Sr61*, possibly because these *Avr* genes were not present in this library which represents about half of the annotated genes encoding secreted proteins expressed in haustoria. Thus, identification of *Avr* gene candidates from four out of seven *Sr* genes screened represents a high detection rate and synthesis of a larger effector library may allow identification of additional candidates.

Specific recognition of the *AvrSr13* and *AvrSr22* candidates by *Sr13c* and *Sr22*, respectively, was confirmed by co-transformation of protoplasts with the individual *Avr* gene candidates and the corresponding *R* genes, which resulted in a substantial reduction in YFP-positive cells compared to the *R* or *Avr* genes alone (Fig. 3a). Similarly, specific recognition of these *Avr* gene candidates was also observed following transformation of protoplasts derived from stable transgenic wheat lines expressing either *Sr13c* or *Sr22*, as well as in protoplasts from native wheat lines expressing *Sr13a* or *Sr22* (Fig. 3b,c). Transient agrobacterium expression assays in *Nicotiana tabacum* and *N. benthamiana* also showed cell death induction upon co-expression of *AvrSr13* with *Sr13c* or *AvrSr22* with *Sr22*, but not with non-matching *R*-*Avr* gene pairs (Fig. 3d,e, Supplementary Fig. 4).

**Fig. 3.**
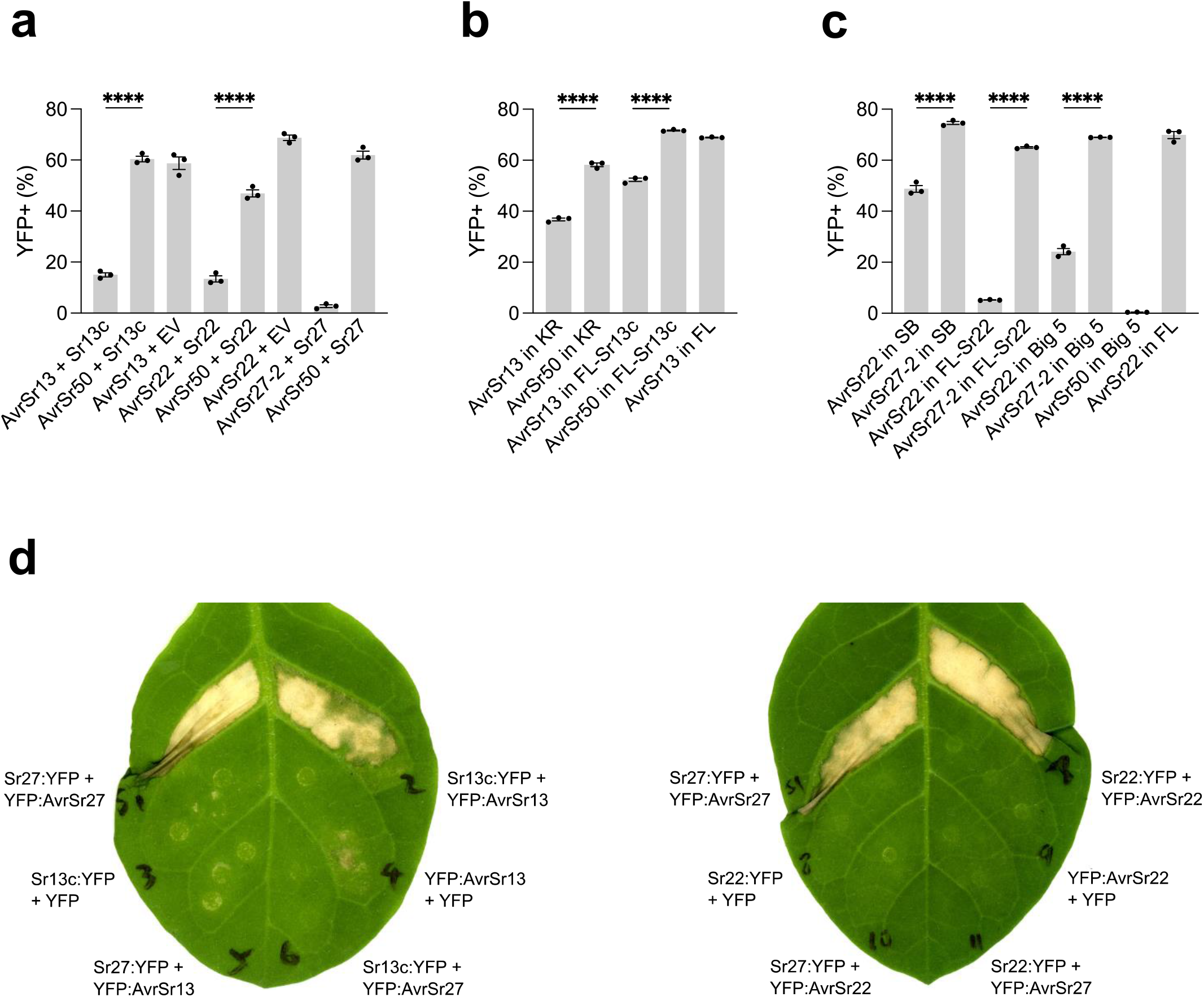
Validation of *AvrSr13* and *AvrSr22* candidates. **a**, Protoplasts were co-transformed with YFP and *AvrSr13, AvrSr22, AvrSr27-2* or *AvrSr50* with an empty vector (EV), *Sr13c, Sr22* or *Sr27*. **b**, YFP was co-transformed with *AvrSr13* or *AvrSr50* into protoplasts from wheat lines Kronos (KR, containing native *Sr13a*), Fielder (FL) or transgenic Fielder containing the *Sr13c* transgene (FL Sr13c) **c**, The YFP reporter was co-transformed with *AvrSr22* or *AvrSr27*-*2* into wheat lines Schomburgk (SB, containing native *Sr22*), Fielder (FL), transgenic Fielder containing the *Sr22* transgene (FL Sr22), or transgenic Fielder containing a five *R* gene cassette including *Sr22* (FL Big5). Plots show the percentage of YFP-positive living cells (mean of three replicates [grey dots] with significant differences indicated; ****, *p* < 0.0001). **d**, *Agrobacterium*-mediated transient co-expression of Sr27, Sr13c or Sr22 (C-terminally fused to YFP) with AvrSr13, AvrSr22, or AvrSr27-2 (N-terminally fused to YFP) or YFP alone in *N. tabacum* leaves.

*AvrSr13* is encoded by PGT21_021053 on chromosome 1A in the Pgt21-0 genome reference, with a null allele on chromosome 1B indicating that this strain is heterozygous for avirulence to *Sr13*. However, the Ug99 *Pgt* isolate (race TTKSK) is homozygous for identical *AvrSr13* genes (PGTUg99_007363 and unannotated) in the A and C haplotypes and is therefore likely homozygous for avirulence on *Sr13. AvrSr22* is encoded by PGT21_017626 on chromosome 16B in the Pgt21-0 genome reference and the alternative allele on chromosome 16A in Pgt21-0 contains a related sequence that is not annotated but encodes a mature protein with nine amino acid differences from *AvrSr22*. Mapping RNA-seq data from Pgt21-0 haustoria and infected plant samples ^22^ confirmed the expression of this transcript during infection at similar levels to *AvrSr22*, suggesting that it is a functional gene which we designated as the *AvrSr22b* allele. The Ug99 genome contains sequences identical to both *AvrSr22* (PGTUg99_032354) and *AvrSr22b* (tig00002160, unannotated).

Co-expression of *AvrSr22b* with *Sr22* in wheat protoplasts resulted in a decrease in the proportion of YFP positive protoplasts (Supplementary Fig. 5a) and also induced cell death in transiently transformed *N. tabacum* and *N. benthamiana* leaves (Supplementary Fig. 5b). Since both *AvrSr22* alleles are recognised by *Sr22*, Pgt21-0 and Ug99 are likely homozygous for avirulence on *Sr22*. The observation that the Ug99 strain is homozygous for avirulence on *Sr13c* and *Sr22* suggests that these resistance genes are more likely to provide durable resistance to Ug99-derived *Pgt* strains than *Sr27, Sr35* or *Sr50* for which this strain is heterozygous for avirulence ^11, 12, 14^.

In summary, we developed a new genetic platform to rapidly identify interacting pairs of plant immunoreceptors and pathogen Avr effectors that confer disease resistance that could be applied to accelerate advances in many critical plant diseases. This will provide molecular tools needed for surveillance of genetic variation at *Avr* loci in pathogen populations and enable the creation of *R-Avr* gene atlases ^23^ to inform breeding and deployment of disease resistant crops. For instance, the homozygosity of the Ug99 stem rust strain for *Avr* genes recognised by *Sr13c* and *Sr22* provides a rationale for prioritising these resistance genes in breeding programs and *R* gene stacking approaches ^3^ targeting durable resistance to this race group. The protoplast library screening platform developed here could also be adapted to identify genes controlling other important biological traits with detectable phenotypes in protoplasts, or that can be linked to cell death/survival or a fluorescence reporter output.

## Supplementary Figure Legends

**Supplementary Fig. 1.**
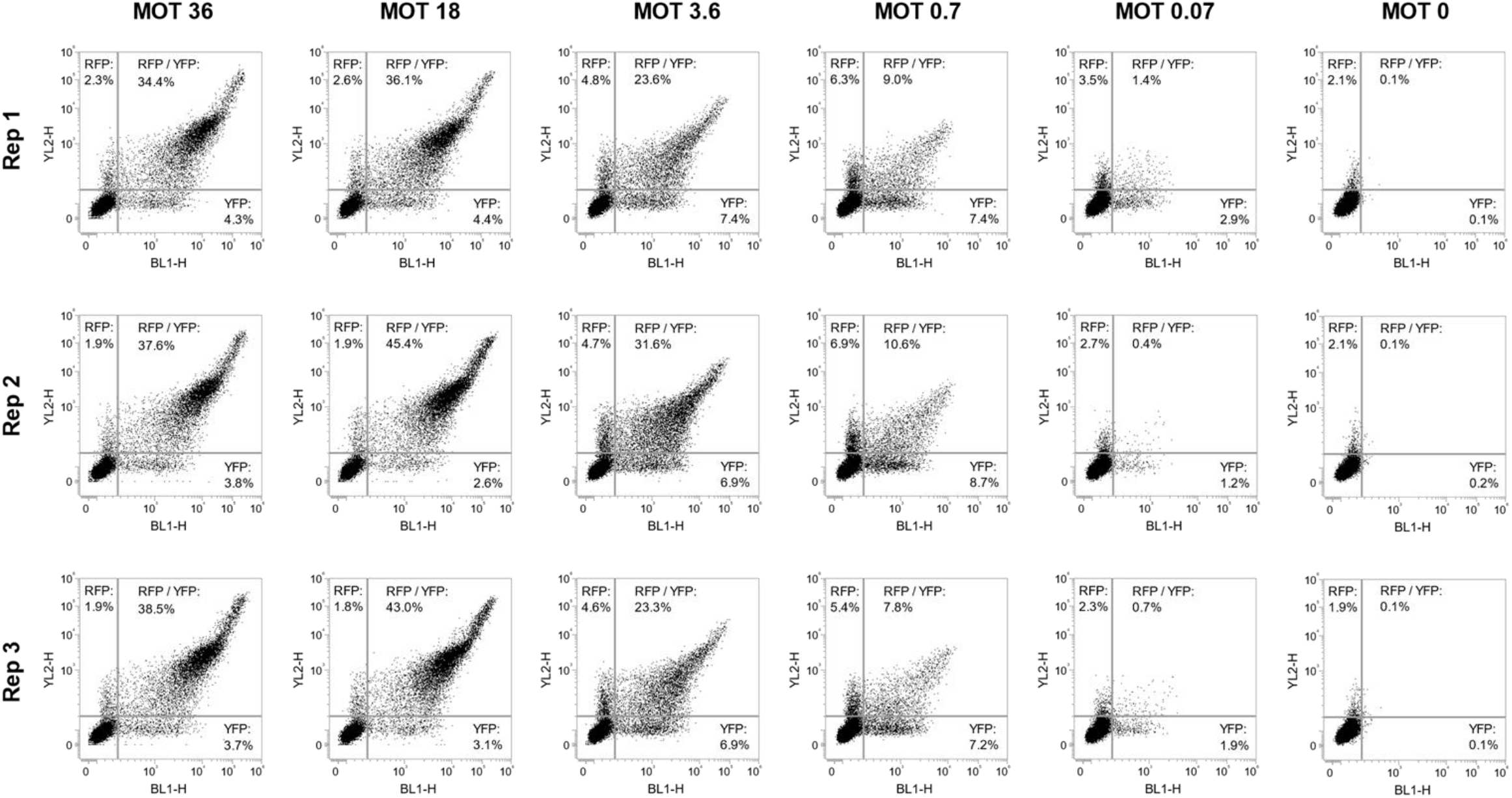
Flow cytometric assessment of single and double construct transformation frequency at different multiplicities of transfection (MOT). The YFP reporter construct (pTA22-YFP) was co-transformed with the RFP reporter construct (pTA22-mRuby3) and the empty vector (pTA22). pTA22-YFP and pTA22-mRuby3 were delivered in equimolar amounts at a various MOT’s (0, 0.07, 0.7, 3.6, 18, 36 million plasmid molecules/cell), while the amount of pTA22 was varied such that the MOT of all constructs combined remained constant at 72 million molecules/cell. Graph shows fluorescence levels of each living cell (dots) for YFP (X-axis log_10_ scale) and RFP (Y-axis log_10_ scale). Gating parameters for YFP and RFP fluorescence are indicated by the grey lines. Data points in the bottom right (rectangular) quadrant represent cells that express YFP only, data points in the top left (rectangular) quadrant represent cells that express RFP only (red), and data points in the top right (square) quadrant represent cells that express both YFP and RFP (yellow).

**Supplementary Fig. 2.**
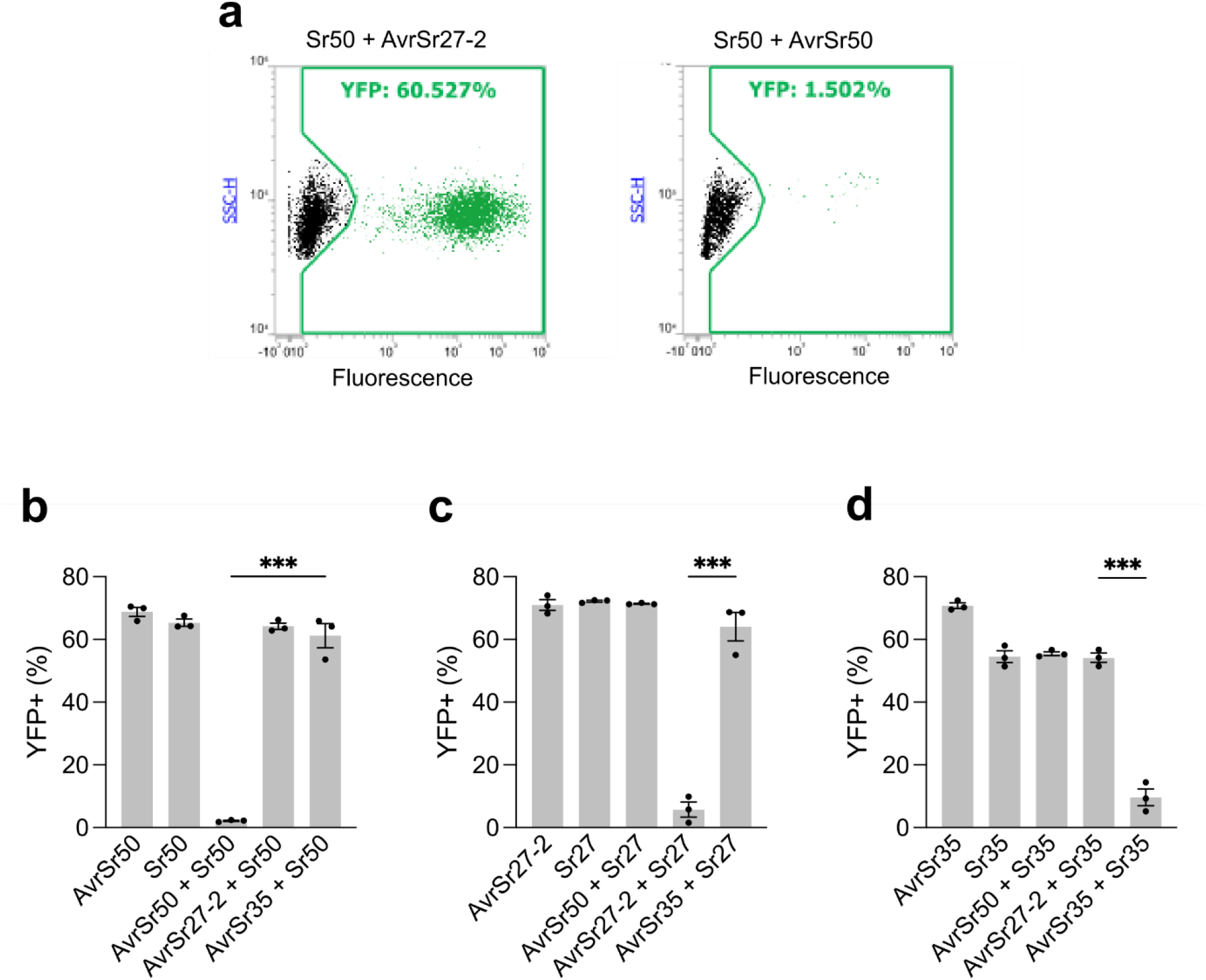
**a**, Sample graphs showing flow cytometry output for protoplasts co-transformed with YFP and Sr50+AvrSr27-2 (top panel) or Sr50+AvrSr50 (bottom panel). YFP fluorescence levels (X-axis log_10_ scale) of each cell (dots) are represented versus side scatter (SSC) (Y-axis log_10_ scale). YFP-positive cells are shown in green with the gating parameters indicated by a green line. **b-d**, Protoplasts were co-transformed with YFP and either single Avr genes (*AvrSr50, AvrSr27-2*, or *AvrSr35*), single R genes (*Sr50, Sr27*, or *Sr35*), matching R-Avr pairs, or non-matching R-Avr pairs. Plots show the percentage of living cells that are YFP-positive (mean of three replicates [grey dots] with significant differences indicated; ****, *p* < 0.0001; ***, *p* < 0.001, two-sample *t*-test equal variances).

**Supplementary Fig. 3.**
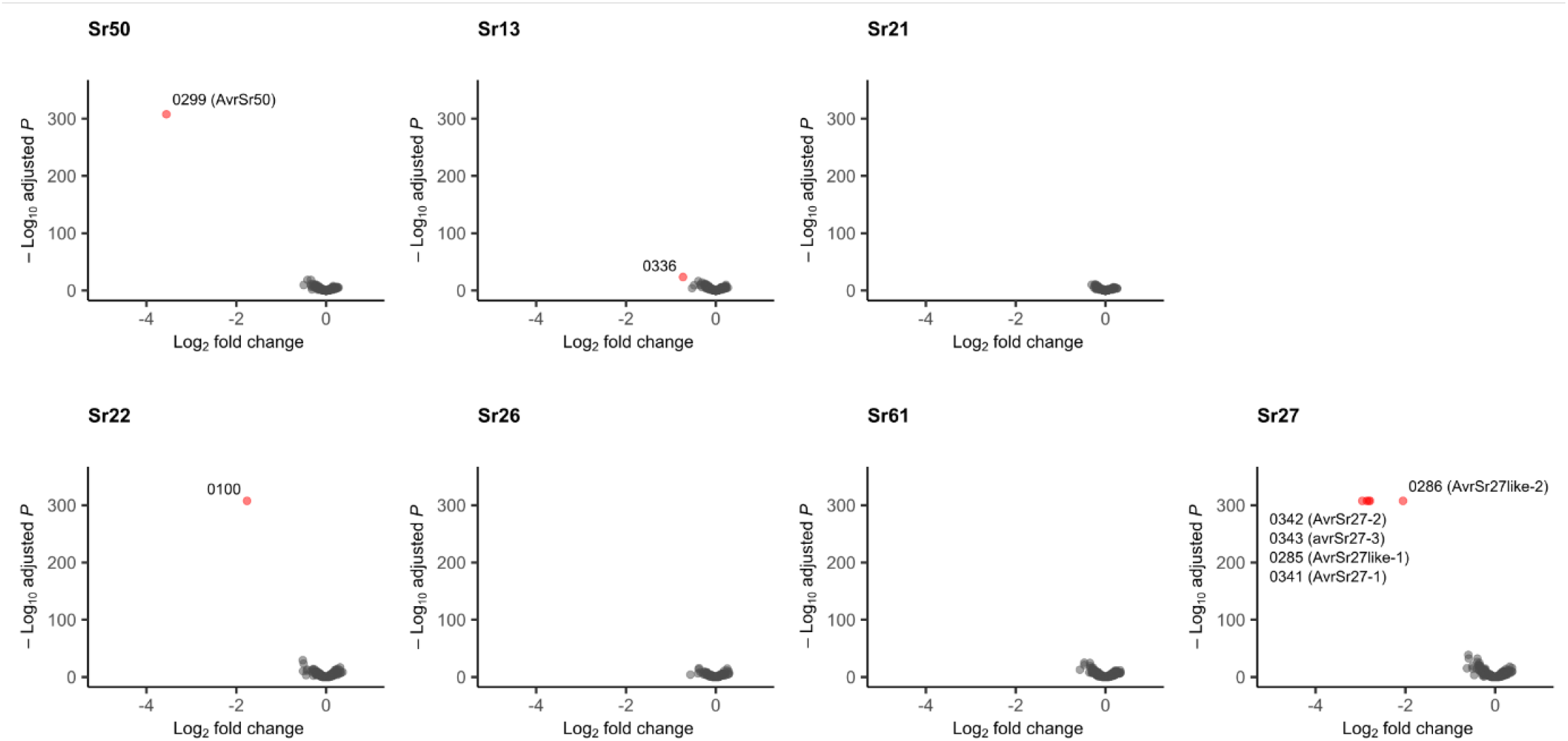
Screening of the pooled stem rust effector library in wheat protoplasts. Expression analysis of the pooled effector library co-transformed into wheat protoplasts with wheat resistance genes Sr50, Sr27, Sr13c, Sr21, Sr22, Sr26 or Sr61. Graph shows a volcano plot of differential expression (log2 scale X-axis) versus adjusted P value (-log10 scale Y-axis) for each of 696 effector constructs. Effectors showing significantly reduced expression in the presence of the R gene compared to the empty vector control are labelled with their library ID number. These data are from an independent screen to that presented in Fig. 1d.

**Supplementary Fig. 4.**
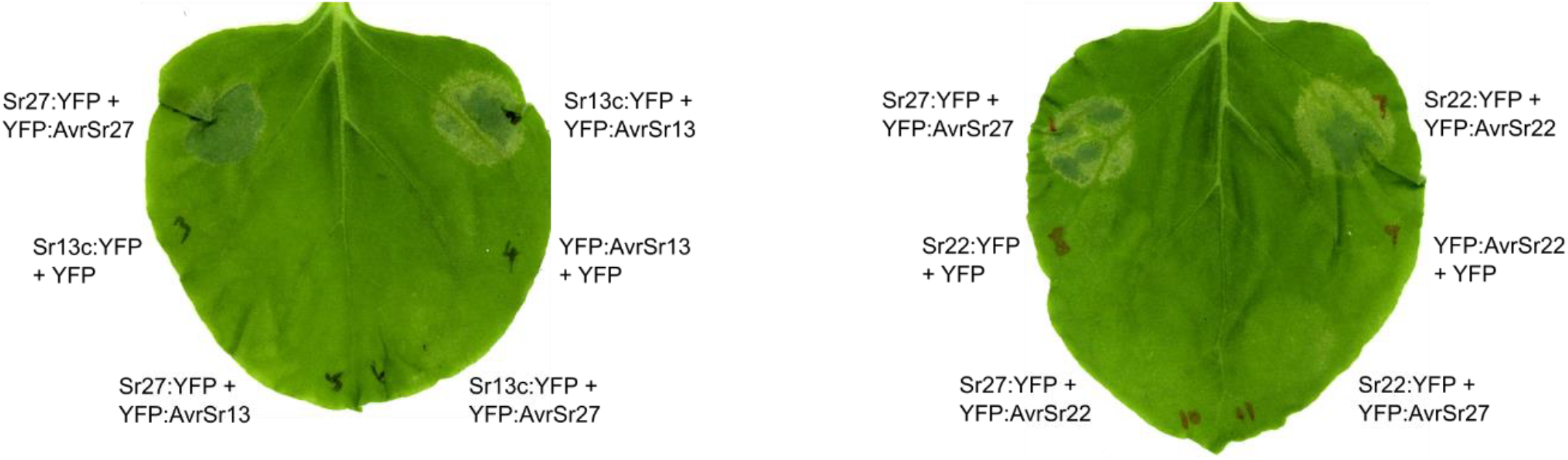
Validation of candidates for AvrSr13 and AvrSr22 via agroinfiltration of *N. benthamiana* leaves. Leaf sectors express combinations of Sr27, Sr13c or Sr22 C-terminally fused to YFP along with AvrSr13, AvrSr22, or AvrSr27 N-terminally fused to YFP, or with YFP alone. Recognition of the Avr protein by the R protein results in cell death, which is visible as discoloured necrotic tissue.

**Supplementary Fig. 5.**
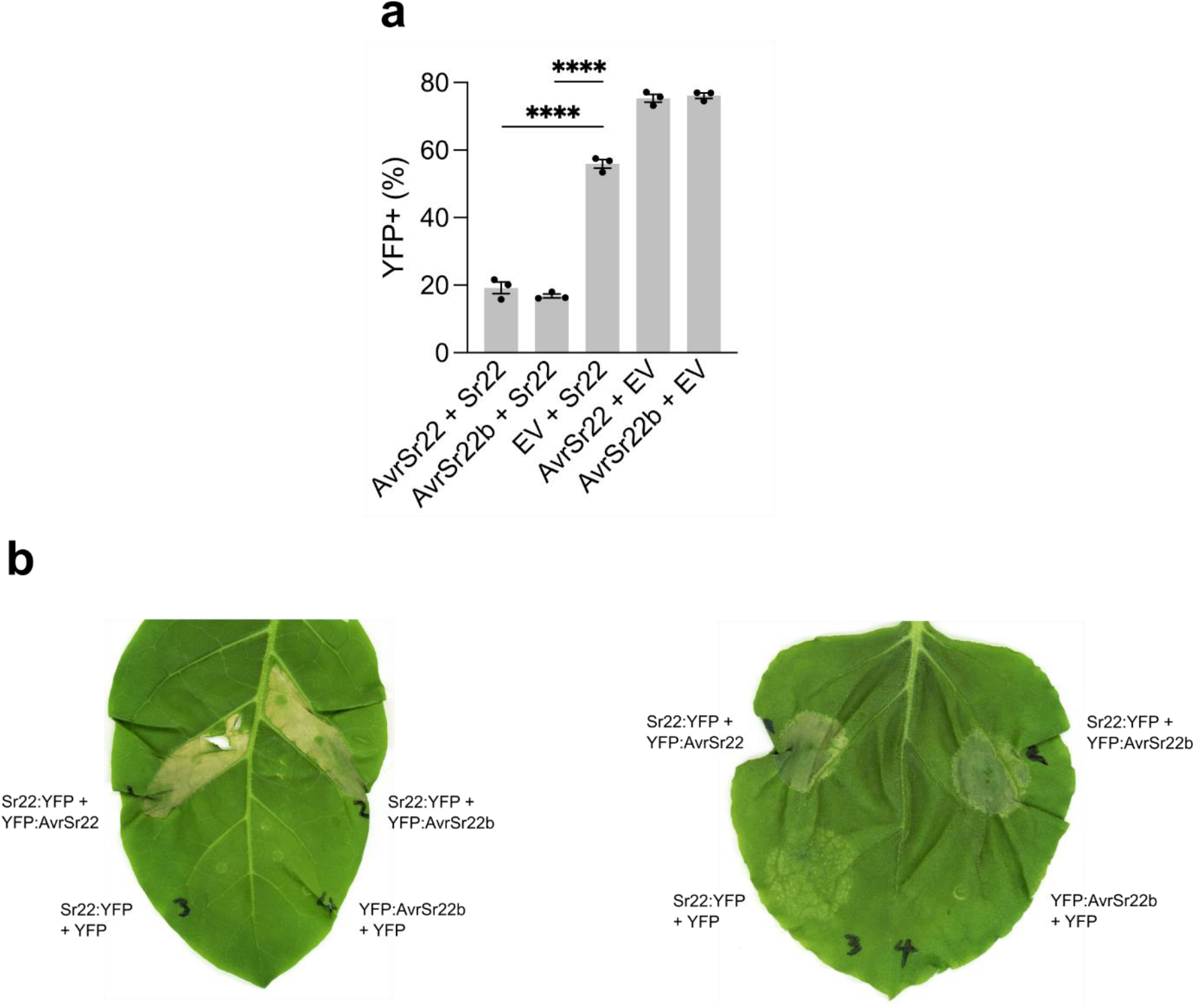
Sr22 recognises both AvrSr22 and AvrSr22b. **a**, The YFP reporter construct was co-transformed with the AvrSr22 (#0100) or AvrSr22b constructs and an empty vector or the Sr22 construct. Plot shows the percentage of living protoplasts that are YFP-positive as the mean of three replicates (grey dots) with significant differences indicated (****, *p* < 0.0001; two-sample *t*-test assuming equal variances). **b**, Agrobacterium-mediated transient expression of AvrSr22 and AvrSr22b with Sr22 in *N. tabacum* (left) or *N. benthamiana* (right) leaves. Image taken 5 days post-infiltration.

**Supplementary Fig. 6.**
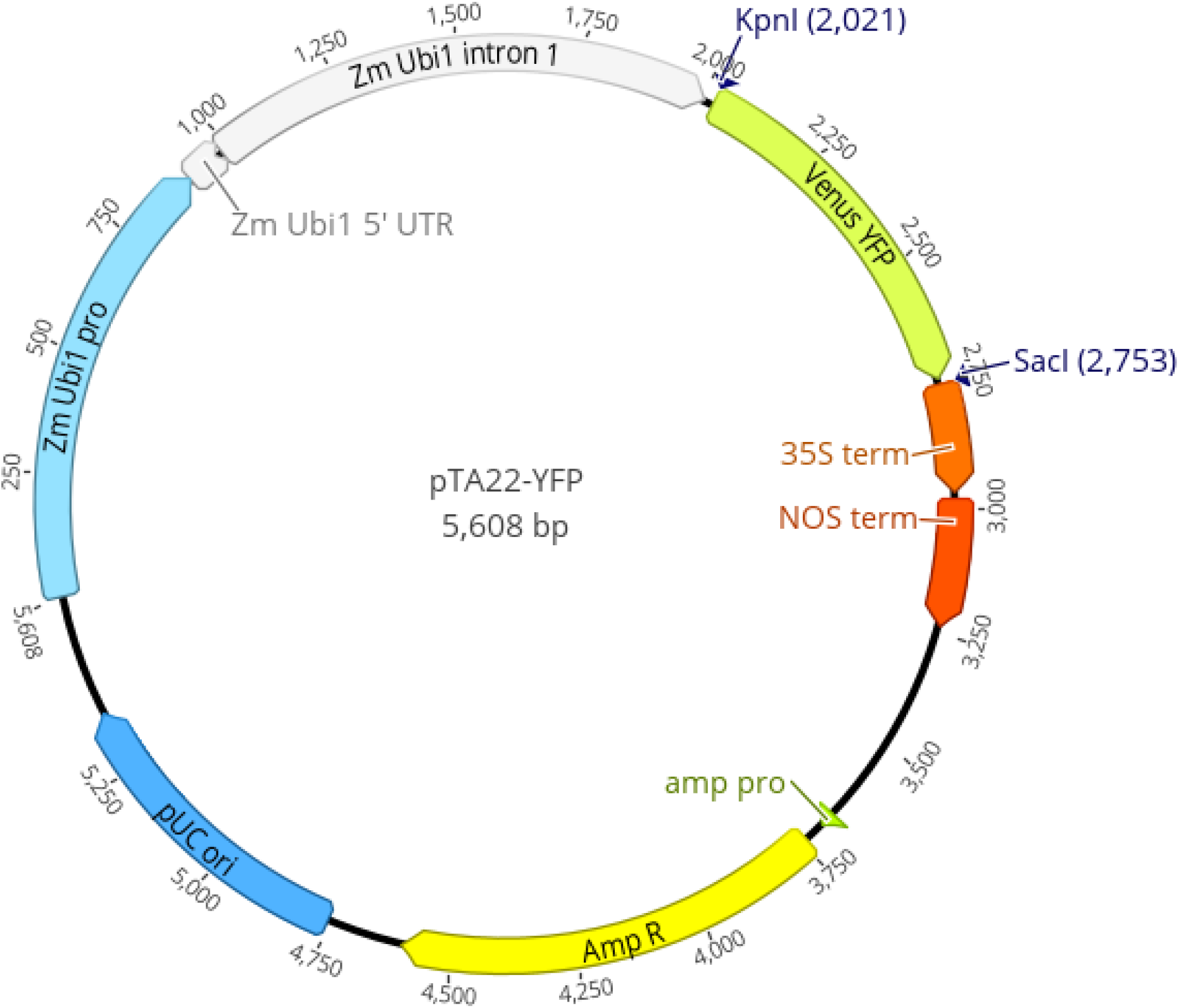
Vector diagram of the YFP reporter construct pTA22-YFP.

## Online Methods

### Vector design and construction

pTA22-YFP (Supplementary Fig. 6) is a high-copy (pUC19 origin of replication) plasmid containing the coding sequence for Venus yellow fluorescent protein (YFP) flanked by the maize ubiquitin 1 promoter (including first intron) and a 35S:NOS double terminator. The negative control empty vector pTA22 was created by digestion of pTA22-YFP with *Kpn*I-*Sac*I followed by blunting and self-ligation, to remove the YFP coding sequence. The destination vectors pTA22-GW and pTA22-GW-PBS were created by replacing the YFP coding sequence in pTA22-YFP with a synthesised Gateway cassette via restriction/ligation (*Kpn*I-*Sac*I). pTA22-GW-PBS is identical to pTA22-GW with the exception that pTA22-GW-PBS also contains a reverse primer binding site (PBS) between the Gateway cassette and the 35S:NOS double terminator.

The coding sequence of the wheat stem rust resistance gene *Sr50* and the open reading frames including introns of the wheat stem rust resistance genes *Sr27, Sr35, Sr13c, Sr21, Sr22, Sr26* and *Sr61* were cloned into pTA22-GW via Gateway LR reaction to create the expression vectors pTA22-Sr50, pTA22-Sr27, pTA22-Sr35, pTA22-Sr13c, pTA22-Sr21, pTA22-Sr22, pTA22-Sr26, and pTA22-Sr61, respectively. The open reading frames in the *R* gene expression vectors were sequence verified. *Avr* gene open reading frames (AvrSr50, AvrSr27-2, AvrSr35, AvrSr22b) were cloned into pTA22-GW-PBS via Gateway LR reaction to create the expression vectors pTA22-AvrSr50-PBS, pTA22-AvrSr27-2-PBS, pTA22-AvrSr35-PBS, and pTA22-AvrSr22b-PBS, respectively. The *Avr* gene expression vectors were confirmed correct by diagnostic restriction enzyme digest.

For transient expression in *Nicotiana* spp, the coding sequences of AvrSr13 and AvrSr22 were transferred from the library constructs pTA22-0336-PBS and pTA22-0100-PBS, respectively, into the pDONR207 vector via Gateway BP reaction (Invitrogen). The predicted coding sequences of Sr13c, Sr22 and AvrSr22b were synthesized and subcloned into pDONR207 via Gateway BP cloning. *R* and *Avr* gene sequences were then transferred into the binary vectors pAM-35s-GWY-YFPv and pBIN19-35s-YFPv-GWY, respectively, by Gateway LR reaction as previously described ^20, 24^.

### Effector library design and construction

Effector candidates from *Pgt* were selected from the genome reference annotation for Pgt21-0 ^11^ (NCBI BioProject PRJNA516922) and a differential expression analysis of all genes encoding secreted proteins that identified eight clusters of genes with different expression profiles during the *Pgt* infection cycle ^12^. Candidates for inclusion in the *Pgt* effector library were selected based on the following filtering criteria: present in secreted protein gene expression clusters 2, 3 or 7; length <1000 nt after removal of the signal peptide encoding region; expression level in haustoria >5 transcripts per million (TPM); SignalP3.0 signal peptide prediction probability >0.5; unique translated protein sequence in candidate set. The coding sequences of the resulting 718 putative effectors were codon-optimised for wheat using GeneOptimizer. Additional sequences designed to facilitate synthesis and cloning were added immediately upstream (5 ‘-AGGCTTCACC-3 ‘) and immediately downstream (5 ‘- CCATACCCAGCTTTCTTGTACAAAGTGGTTTGATCGTACTGTCGAGATCTAGCA*ACGCGATCGGAGGCGCTCAT TATACGCAGATTCTTTATCGAAGCTGAGGGTGTGCCCGCTGTAACCCGCAAAGCCGTCAATATACAATCCTGACCAAATAGGAGACTGAACCGGTTTGGTAGCAGATAAGTTGCTTGGTGCCG*-3 ‘) of the optimised coding sequences. It was a manufacturing requirement that all synthesised fragments be at least 300 bp long. Therefore, the minimum length of randomly generated filler sequence (italicised above in the downstream sequence) was used where necessary (the 94 smallest effectors) to meet this minimum length. Six hundred and ninety-six of the 718 putative effectors were successfully synthesised and cloned (Twist Bioscience) into pTA22-GW-PBS to create a library of expression vectors with names pTA22-0001-PBS to pTA22-0718-PBS.

### Effector library pooling and propagation

The 696 effector library constructs were individually resuspended in 100 µL IDTE buffer (10 mM Tris, 0.1 mM EDTA, pH 7.5) and then pooled in equimolar amounts (150 fmol per construct) using a JANUS G3 automated liquid handling workstation. To propagate the pooled library, 2.5 µL of pooled library DNA (∼330 ng total) was transformed into 50 µL of ElectroMAX Stbl4 competent *Escherichia coli* cells (Invitrogen) by electroporation, followed by addition of 2.5 mL pre-warmed SOC medium (Invitrogen). Outgrowth without selection proceeded at 37°C for 1 hour with shaking at 220 RPM. After outgrowth, cells were spun down and resuspended in 2.5 mL of Luria-bertani (LB) medium. The culture was then diluted 1 in 200 using LB medium, and 300 µL of the diluted culture was spread on a 14.5 cm diameter LB agar plate containing 50 µg/mL carbenicillin. Colonies were scraped off the plates using a cell spreader and LB medium, and then transferred to Falcon tubes. Cells were spun down, supernatant was removed, and the pellets were then stored at -20°C prior to plasmid DNA isolation.

### Plasmid DNA isolation

pTA22-YFP was isolated using the MACHEREY NAGEL NucleoBond Xtra Maxi Plus EF kit. All other plasmids were isolated using the MACHEREY NAGEL NucleoBond Xtra Midi Plus EF kit. The pooled library was isolated using the QIAGEN EndoFree Plasmid Giga kit. Isolated plasmid DNA was resuspended at a concentration of 1 µg/uL (measured on NanoDrop).

### Protoplast isolation and transformation

Wheat (*Triticum aestivum*) seeds were planted in 13 cm pots (12 seeds per pot) containing Martins Seed Raising and Cutting Mix supplemented with 3 g/L osmocote. Seedlings were grown in a growth cabinet at 24°C on a cycle of 12 hours light (∼100 µmol m^-2^ s^-1^) and 12 h dark, for 7–8 days. Wheat cultivar Fielder was used unless otherwise stated. Protoplast isolation and transformation was carried out as described previously ^25^, except that released protoplasts were filtered through a 40 μm nylon cell strainer and final suspension was in 6 mL MMG solution (4 mM MES-KOH (pH 5.7), 0.4 M mannitol, 15 mM MgCl_2_). The protoplast concentration was determined by cell counting on a hemocytometer, and subsequently adjusted to 2.5 × 10^5^ cells/mL using MMG solution. For standard individual transformations involving single effector genes, 3 pmol of each vector was mixed with 200 μL of protoplasts (50,000 protoplasts) and 230 μL of PEG solution (40% (wt/vol) polyethylene glycol-4000, 0.2 M mannitol, 100 mM CaCl_2_) in a 2 mL tube. The DNA/protoplast/PEG mixture was homogenised by gently flicking the tube, and then incubated for 15-30 min at room temperature before adding 940 μL of W5 solution and gently inverting the tube to mix. Transformed protoplasts were centrifuged for 2 min at 100 x g and the supernatant was removed. Protoplasts were resuspended in 650 μL W5 solution, transferred to 12-well cell culture plates and incubated at 23°C for 24 hours in the dark. For mock library screening and pooled library screening, transformation reactions were scaled up 8-10x in 25 mL tubes, and transformed protoplasts were incubated in cell culture flasks. Transformations were performed in triplicate for all treatments and controls.

### Flow cytometry

After the 24-hour incubation period, protoplasts were transferred to 2 mL tubes, stained with propidium iodide (10 µg/mL) and then subjected to flow cytometry using the Invitrogen Attune NxT Flow Cytometer to detect fluorescence properties of individual cells. Propidium iodide fluorescence (excitation 561 nm, emission filter 620/15 nm) was measured as an indicator for living protoplasts (no fluorescence), and YFP fluorescence (excitation 488 nm, emission filter 530/30 nm) was measured as an indicator of the reporter gene expression. The percentage of YFP-positive protoplasts in the living (propidium iodide-negative) population was used to assess the transformation efficiency and the strength of R-Avr induced PCD. When RFP was used as a reporter, cells were not stained with propidium iodide and RFP fluorescence was detected using excitation 561 nm, emission filter 620/15 nm.

### mRNA extraction and cDNA synthesis/PCR

After the 24-hour incubation period, protoplasts were transferred to 5 mL tubes and centrifuged for 3 min at 150 x g. The supernatant was discarded, and mRNA was extracted from the protoplast pellet using the Invitrogen Dynabeads mRNA DIRECT Purification Kit, according to the manufacturer‘s protocol for ‘Mini’ extractions. mRNA was eluted with 20 µL of elution buffer and the concentration determined using the Invitrogen Qubit RNA HS Assay Kit with the Invitrogen Qubit 4 Fluorometer. Concentrations ranged from ∼5-9 ng/µL. Library-specific cDNA synthesis and PCR was carried out using the Invitrogen SuperScript IV One-Step RT-PCR System with ezDNase kit, following the manufacturer’s protocol with minor modifications. All reactions used 30 ng of mRNA template. The forward primer ZmUbi1_5UTR_F3b (GCACACACACACAACCAG) was used with the reverse primer FS_cDNA_R (TGCTAGATCTCGACAGTACG). Cycling conditions were as follows: 55°C for 10 min (inactivation of ezDNase and first strand cDNA synthesis), 98°C for 2 min (inactivation of reverse transcriptase and initial denaturation), 98°C for 10 sec (denaturation), 61°C for 10 sec (annealing), 72°C for 35 sec (extension), 72°C for 5 min (final extension). Eighteen cycles of denaturation, annealing and extension were performed. The PCR product was column purified using the QIAGEN QIAquick PCR Purification Kit, following the manufacturer’s protocol. DNA was eluted with 35 µL of elution buffer and the concentration determined using the Qubit 1X dsDNA HS Assay Kit with the Invitrogen Qubit 4 Fluorometer. Concentrations ranged from ∼8-21 ng/µL.

### Illumina library construction and RNA-Seq

Illumina libraries were constructed using the Illumina DNA Prep Kit and IDT for Illumina DNA/RNA UD Indexes, following the manufacturer’s protocol with minor modifications. Around 120-220 ng of dsDNA from the cDNA synthesis/PCR was used as input to achieve on-bead normalisation. Right side library cleanup with purification beads was carried out as per the Illumina protocol, while a 1.8x bead ratio was used for the left side clean up in order to retain smaller amplicons. Illumina library concentrations were measured using the Qubit 1X dsDNA HS Assay Kit with the Invitrogen Qubit 4 Fluorometer. Concentrations ranged from ∼22-29 ng/µL. Quality control of library size was carried out using the Agilent TapeStation 2200 with High Sensitivity D1000 ScreenTapes and Reagents. Illumina libraries were pooled in equimolar amounts for RNA-seq. Pooled Illumina libraries were sequenced on the Illumina NextSeq 500 platform (ACRF Biomolecular Resource Facility, The John Curtin School of Medical Research, Australian National University) using the NextSeq 500/550 Mid Output Kit v2.5 and 74 bp paired end reads. PhiX was spiked in at 5%.

### Analysis of differential gene expression

RNA sequencing reads were cleaned using fastp 0.22.0 ^26^ (--length_required 20) specifying the UTR sequences common to all library transcripts as well as the Illumina DNA Prep adapter sequence as adapters. The clean reads were aligned to the coding sequences of the 696 cloned effector candidates with HISAT2 2.2.1 ^27^ (--very-sensitive; --sp 1,1; --no-spliced-alignment). Mappings where the read pairs map to different transcripts were dismissed. Salmon 1.8.0 ^28^ was used to quantify expression from the HISAT2 alignments. Read counts were imported into DESeq2 (Love et al., 2014) with tximport (type = “salmon “). Differential expression analysis was performed with DESeq2 ^29^ and default parameters followed by lfcShrink (type= “apeglm “). P-values of zero (-Log_10_ adjusted P = infinity) were converted to the machine-lowest value possible in R (function: .Machine$double.xmin) resulting in a -Log_10_ adjusted P value slightly greater than 300 for those data points (AvrSr50 in both screens; 0100 and the five variants of AvrSr27 in screen 2). Volcano plots were produced with EnhancedVolcano (https://github.com/kevinblighe/EnhancedVolcano). For the mock library screen, AvrSr50 and AvrSr27-2 transcripts per million (TPM) was calculated from the read counts from the HISAT2 alignments and normalised to the AvrSr35 TPM. Significance was assessed with an unpaired *t*-test.

### Agroinfiltration of *Nicotiana tabacum* and *Nicotiana benthamiana* leaves

*N. tabacum* and *N. benthamiana* plants were grown in a growth chamber at 23 °C with a 16-hour light period. *Agrobacterium tumefaciens* cultures containing the expression vectors of each construct were grown overnight at 28°C in LB media with appropriate antibiotic selections. The cells were pelleted and resuspended in infiltration mix (10 mM MES pH 5.6, 10 mM MgCl_2_, 1500 μM acetosyringone) to an optical density (OD600) of 0.2 or 0.5, followed by incubation at room temperature for two hours. Cultures were infiltrated into leaves of four-week-old plants with a 1 mL syringe. For documentation of cell death, leaves were photographed or scanned 2–5 days after infiltration.

## Acknowledgements

This work was supported by the CSIRO Synthetic Biology Future Science Platform and the CSIRO Research Office.

---

**Supplementary Table S1.** plasmids used in this study

| Use | Construct | Plasmid name | Plasmid backbone | Cloning method | Source |
| --- | --- | --- | --- | --- | --- |
| Entry vectors for cloning | AvrSr13 | pDONR-AvrSr13 | pDONR207 | BP-Gateway | - |
|  | AvrSr22 | pDONR-AvrSr22 | pDONR207 | BP-Gateway | - |
|  | AvrSr22b | pDONR-AvrSr22b | pDONR207 | synthesis/BP-Gateway | - |
|  | Sr13c | pDONR-Sr13c | pDONR207 | synthesis/BP-Gateway | - |
|  | Sr22 | pDONR-Sr22 | pDONR207 | synthesis/BP-Gateway | - |
| Destination vectors for cloning | Gateway (attR1 / attR2) | pTA22-GW | pTA22 | restriction / ligation | - |
|  | Gateway (attR1 / attR2) | pTA22-GW-PBS | pTA22 | restriction / ligation | - |
| Expression vectors for wheat protoplasts | Ubi1p:Venus:35St:NOST | pTA22-YFP | pTA22 | restriction / ligation | - |
|  | Ubi1p:mRuby3:35St:NOST | pTA22-mRuby3 | pTA22 | restriction / ligation | - |
|  | Ubi1p-35St:NOST (empty) | pTA22 | pUC19-like | restriction / ligation | - |
|  | Ubi1p:Sr13c:35St:NOST | pTA22-Sr13c | pTA22-GW | LR-Gateway | - |
|  | Ubi1p:Sr21:35St:NOST | pTA22-Sr21 | pTA22-GW | LR-Gateway | - |
|  | Ubi1p:Sr22:35St:NOST | pTA22-Sr22 | pTA22-GW | LR-Gateway | - |
|  | Ubi1p:Sr26:35St:NOST | pTA22-Sr26 | pTA22-GW | LR-Gateway | - |
|  | Ubi1p:Sr61:35St:NOST | pTA22-Sr61 | pTA22-GW | LR-Gateway | - |
|  | Ubi1p:Sr50:35St:NOST | pTA22-Sr50 | pTA22-GW | LR-Gateway | - |
|  | Ubi1p:Sr27:35St:NOST | pTA22-Sr27 | pTA22-GW | LR-Gateway | - |
|  | Ubi1p:Sr35:35St:NOST | pTA22-Sr35 | pTA22-GW | LR-Gateway | - |
|  | Ubi1p:AvrSr13:35St:NOST | pTA22-0336-PBS | pTA22-GW-PBS | synthesis | - |
|  | Ubi1p:AvrSr22:35St:NOST | pTA22-0100-PBS | pTA22-GW-PBS | synthesis | - |
|  | Ubi1p:AvrSr22b:35St:NOST | pTA22-AvrSr22b-PBS | pTA22-GW-PBS | LR-Gateway | - |
|  | Ubi1p:AvrSr50:35St:NOST | pTA22-AvrSr50-PBS | pTA22-GW-PBS | LR-Gateway | - |
|  | Ubi1p:AvrSr27-2:35St:NOST | pTA22-AvrSr27-2-PBS | pTA22-GW-PBS | LR-Gateway | - |
|  | Ubi1p:AvrSr35:35St:NOST | pTA22-AvrSr35-PBS | pTA22-GW-PBS | LR-Gateway | - |
| Expression vectors in *Nicotiana* | 35Sp:AvrSr27-2:35St | pBIN19-YFP-AvrSr27-2 | pBIN19-YFP-GTW | LR-Gateway | Upadhyaya et al 2021 |
|  | 35Sp:Sr13c-YFP:35St | pAM-Sr13c-YFP | pAM-GTW-YFP | LR-Gateway | - |
|  | 35Sp:Sr22-YFP:35St | pAM-Sr22-YFP | pAM-GTW-YFP | LR-Gateway | - |
|  | 35Sp:YFP-AvrSr13:35St | pBIN19-YFP-AvrSr13 | pBIN19-YFP-GTW | LR-Gateway | - |
|  | 35Sp:YFP-AvrSr22:35St | pBIN19-YFP-AvrSr22 | pBIN19-YFP-GTW | LR-Gateway | - |
|  | 35Sp:YFP-AvrSr22b:35St | pBIN19-YFP-AvrSr22b | pBIN19-YFP-GTW | LR-Gateway | - |
|  | 35Sp:Sr27-YFP:35St | pAM-Sr27-YFP | pAM-GTW-YFP | LR-Gateway | Upadhyaya et al 2021 |

